# Late radiation-induced brain injury is associated with functional cerebrovascular impairment without structural vessel loss in mice

**DOI:** 10.64898/2026.11.03.763950

**Authors:** Michelle R. Tamplin, Anthony H. Vitale, Pedro Loureiro, Anne Cunningham, Chu-Yu Lee, Eric D. Axelson, Vikram Subramanian, Li-Hsien Lin, William Thiel, Michael S. Petronek, Isabella M. Grumbach

## Abstract

**Background:** Radiation-induced brain injury (RIBI) is a major and often irreversible complication of cranial radiotherapy, affecting up to 90% of long-term survivors. Although cerebrovascular damage is a leading mechanistic hypothesis, whether chronic resting hypoperfusion occurs independently of structural vessel loss remains unknown. The aim of this study was to comprehensively characterize cerebrovascular, structural, and behavioral outcomes in a murine model of late RIBI to determine the primary drivers of vascular dysfunction.

**Methods:** Adult C57BL/6 mice underwent 12 Gy whole-brain irradiation and were assessed one year later using laser speckle contrast imaging (LSCI), 7T MRI, immunohistochemistry, and behavioral testing.

**Results:** Irradiated mice exhibited reduced pial flow velocity and flow area by LSCI, indicative of upstream cerebrovascular dysfunction, while bulk cortical tissue perfusion was preserved. This pial hypoperfusion was accompanied by cerebral atrophy, decreased T2* relaxation time and tissue iron deposition consistent with chronic microhemorrhage across multiple brain regions by MRI, blood-brain barrier disruption reflected by reduced ZO-1 and VE-cadherin expression, and increased anxiety-like behavior. Strikingly, capillary density, pericyte coverage, astrocyte endfeet, and NeuN-positive neuronal nuclei were all preserved.

**Conclusion:** These findings demonstrate that functional vascular impairment - not structural vessel or neuronal loss - drives hypoperfusion and blood-brain barrier disruption in late RIBI. This dissociation between vascular dysfunction and rarefaction reframes the pathophysiology of RIBI and identifies endothelial dysfunction as a tractable therapeutic target for intervention.

## INTRODUCTION

Radiation therapy is a cornerstone of brain cancer treatment, yet its benefits come at significant cost to healthy brain tissue. Between 50–90% of patients surviving 6 months or more after cranial radiotherapy develop irreversible cognitive impairments, including deficits in short-term memory, verbal processing, and executive function, which represent the delayed-onset stages of radiation-induced brain injury (RIBI).^1,2^ These deficits arise after both whole- and partial-brain irradiation due to off-target exposure of healthy neural tissue,^3^ and remain a major unresolved obstacle to survivorship care. The mechanisms driving delayed-onset RIBI are incompletely understood, limiting the development of effective preventative and therapeutic strategies.

Cerebrovascular injury has emerged as a leading mechanistic hypothesis for RIBI. Histological studies have identified early vascular changes, including vessel lumen dilation, endothelial cell nuclear enlargement, and perivascular astrocyte hypertrophy, preceding demyelination and white matter necrosis,^4^ and cognitive impairment has been observed in multiple preclinical models without detectable loss of neuronal structures.^5,6^ Capillary rarefaction has also been proposed as a driver of cognitive decline,^7^ but its occurrence and severity vary substantially across the literature. A recent systematic review of 72 preclinical studies found that cerebrovascular damage is dose-, dose rate-, and field size-dependent, with studies using moderate, clinically-relevant doses reporting preserved vascular structure despite cognitive and vascular impairments.^2^ This raises the possibility that functional vascular impairment, rather than overt vessel loss, may be sufficient to drive RIBI. Consistent with this, neurovascular uncoupling and endothelial dysfunction have been identified as early and persistent features of RIBI that impair stimulus-evoked cerebral blood flow responses.^8,9^ However, whether resting perfusion is chronically reduced after WBI in the absence of capillary rarefaction, and what structural and behavioral consequences accompany such dysfunction, has not been directly addressed.

Prior studies in WBI models have focused on stimulus-evoked blood flow responses, demonstrating early impairment of neurovascular coupling and endothelial reactivity.^9,10^ Resting perfusion, which may more sensitively reflect chronic endothelial dysfunction and baseline metabolic supply, has been largely overlooked as an outcome. Notably, our group recently demonstrated reduced resting retinal perfusion by laser speckle flowgraphy within months of ocular radiotherapy in patients, preceding detectable capillary dropout or vision loss by years.^11^ This clinical observation suggests that hypoperfusion may be an early and sensitive marker of radiation-induced vascular injury, detectable before structural changes emerge. We therefore reasoned that LSCI, a widely available preclinical analog to laser speckle flowgraphy, was well-suited to test whether an analogous dissociation between resting hypoperfusion and vessel loss occurs in the irradiated brain.

Here, we report that 12 Gy WBI in adult mice produces, at one year, a syndrome of pial hypoperfusion, blood-brain barrier disruption, cerebral atrophy, and anxiety-like behavior in the absence of capillary rarefaction or neuronal loss. These findings reframe the vascular pathology of late RIBI, implicating endothelial dysfunction rather than structural vessel loss as the primary driver, and identify the cerebral vasculature as a tractable therapeutic target.

## MATERIALS AND METHODS

### Animal care and irradiation procedure

All experimental procedures complied with the Institute of Laboratory Animal Resources, National Academy of Sciences, and were approved by the University of Iowa and Iowa City VA Health Care System Institutional Animal Care and Use Committees (#9122263 and #2490305). Male C57BL/6J mice (Jackson Laboratories, strain #000664) were housed under controlled temperature (23°C) with a 12-hour light/dark cycle, with free access to water and standard rodent chow. At 10 weeks of age, mice underwent whole-brain irradiation (WBI) or sham treatment using an XStrahl Small Animal Radiation Research Platform (SARRP) at the University of Iowa Radiation and Free Radical Research Core, under anesthesia with 2% isoflurane in 3% oxygen. A mean dose of 12 Gy was prescribed to a single isocenter at the approximate center of the brain using a multivariate collimation system with a 10 mm × 12 mm field size (**Supplementary Fig. 1A,B**). Dose delivery was confirmed by cone beam CT and brain volume contouring in MuriPlan treatment planning software (v.3.0.0; **Supplementary Fig. 1C**). Sham-treated mice underwent identical procedures without irradiation.

At the study endpoint, mice were euthanized by isoflurane overdose (5%) and transcardiac perfusion with PBS followed by 4% paraformaldehyde. Brains were harvested, stored overnight in 30% sucrose at 4°C, cryoembedded in OCT (Tissue-Tek), and sectioned coronally at 10 µm. Slides were stored at −80°C until use.

### Behavioral testing

Mice were acclimated to the testing room and handled by the test operator for three days prior to testing. Anxiety-like behavior was assessed using the open field test, in which mice freely explored a five-sided acrylic arena (40 × 40 cm²) for 10 minutes. Trials were video recorded and analyzed using EthoVision tracking software (v.16, Noldus). The arena was divided into a center zone (20 × 20 cm²) and a border zone; anxiety-like behavior was quantified as the percentage of time spent in the border zone. Locomotor activity was assessed by mean velocity and total distance traveled.

Short-term spatial memory was evaluated using the Y-maze test in a three-arm acrylic arena. In the first trial, mice were placed in the Start arm with the Novel arm blocked by an acrylic insert, and allowed to explore the Start and Familiar arms freely for 5 minutes before returning to the home cage. After 4 hours, mice were re-tested with access to all three arms. Novelty preference was quantified as the percentage of entries into the Novel arm and by the Discrimination Index (DI), defined as:

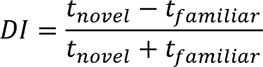

where *t_novel_* and *t_familiar_* are the time spent exploring the Novel and Familiar arms, respectively. All trials were video recorded and tracked in EthoVision.

### Magnetic resonance imaging (MRI)

Mice were imaged on a 7T Discovery MR901 system (GE Healthcare, Milwaukee, WI) using a body transmit coil and 2-channel mouse brain receiver coil to generate anatomical T2-weighted images and T2* maps.¹² Mice were maintained under isoflurane anesthesia (0.5–3% in 100% O₂) throughout. A high-order B0 shimming routine was applied prior to acquisition to minimize field inhomogeneities. Anatomical T2-weighted images were acquired using a 3D FIESTA sequence (in-plane resolution of 104 µm², slice thickness of 160 µm, pixel bandwidth of 326 Hz, flip angle of 30°, TE/TR of 3/6.1 ms, 4 averages, scan time 9 min 28 s). T2* maps were generated from a 2D multi-echo gradient echo (GRE) sequence (in-plane resolution of 156 µm², slice thickness of 500 µm, 18 axial slices, pixel bandwidth of 244 Hz, flip angle of 60°, TR of 1000 ms, 6 TEs from 2.5–22.5 ms in 4 ms increments, 2 averages, scan time 4 min 24 s) using a two-stage fitting procedure applied to GRE magnitude images, as previously described.^12^

Brain regions were defined using DSURQE Mouse Brain Atlas structural labels^13^ registered to individual anatomical images via nonlinear co-registration in Advanced Normalization Tools (ANTs).^14^ T2* maps were then linearly co-registered to the anatomical images labelled with atlas structures. Volume and T2* relaxation time were analyzed in global compartments (total grey matter, total white matter, whole brain) and regions associated with anxiety-like behavior (amygdala, brainstem, cerebrum, cerebellum, frontal cortex, hippocampus, hypothalamus, and periaqueductal gray). A complementary region-based analysis restricted to compartments showing changes in both volume and T2* has been reported separately.^15^

### Laser speckle contrast imaging (LSCI)

Resting cerebral blood flow was measured in anesthetized mice (2–3% isoflurane in 100% O₂) using a laser speckle contrast imaging device (Moor Instruments Ltd). Following a midline incision to expose the skull, ultrasound gel was applied to the skull surface to prevent drying. Blood flow images were acquired at 25 frames per second over 60 seconds, after which mice were euthanized by isoflurane overdose and transcardiac perfusion.

Images were analyzed using moorFLPI Review software (v5.0, Moor Instruments Ltd) by an operator blinded to treatment group; images with artifacts precluding reliable measurement were excluded. A region of interest (ROI) encompassing the cortex between the inferior cerebral vein and transverse sinus was drawn manually. A lower threshold of MBR 500 was applied to exclude artifacts such as hair or bubbles in the image field. Within each ROI, a threshold of MBR 890.9, determined from the flow histogram generated by the analysis software (22 bins, MBR range 200–4000), was used to segment pial vessel flow (above threshold) from cortical tissue flow (below threshold). Mean flux in each compartment was calculated using a custom Python script.

### Protein isolation and immunoblotting

Cerebral microvessels were isolated from flash-frozen brains following isoflurane overdose using an established protocol described in detail elsewhere.^16^ For protein extraction, microvessel pellets were resuspended in 130 µL of RIPA lysis buffer (Fisher Scientific, #R0278) supplemented with protease and phosphatase inhibitors (Pierce, #A32963 and A32957) and agitated for 30 minutes at 15-20°C. Samples were heat-denatured at 95°C for 10 minutes, then sheared using a Covaris E220 focused ultrasonicator (peak incident power 175 W, duty factor 10%, 200 cycles per burst, 300 seconds, 6°C). Lysates were clarified by centrifugation at 16,100 x *g* for 30 minutes at 4°C, and supernatants were stored at -80°C. Protein concentration was determined by BCA assay (Pierce, #23225).

For western blotting, equal protein amounts (10 µg) were resolved on NuPAGE 4–12% Bis-Tris gels (Life Technologies) and transferred to PVDF membranes (Mini-PROTEAN 3, Bio-Rad). Membranes were stained with Ponceau S (Cell Signaling #59803S) to confirm transfer and serve as a loading control, then blocked with 3% BSA or Starting Block™ Buffer (Thermo Fisher #37538) for 2 hours. Primary antibodies (anti-ZO-1 (Millipore, AB2272) and anti-VE-cadherin (BiCell Scientific, 00105), both at 1:1000) were applied overnight at 4°C or for 2 hours at room temperature. After washing, secondary antibodies (1:5000, 2 hours, room temperature) were applied and membranes were developed with Amersham ECL Prime reagent (Cytiva). Densitometry was performed in ImageJ (v.1.54p)^17^ with values normalized to Ponceau S total protein.

### Immunohistochemistry and histology

For neuronal nuclei staining, slides were blocked with 10% donkey serum (1 hour, 25°C), then incubated overnight at 25°C with anti-NeuN (NOVUS, NBP3-05554; 1:200) in blocking buffer. Slides were washed in PBS and incubated with secondary antibody (Jackson, AB 2340422; 1:200) and TOPRO-3 (Invitrogen, T3605; 1:2000) for 2 hours at 25°C, followed by overnight at 4°C. Slides were washed, mounted with ProLong Diamond Antifade Mountant (Invitrogen, P36961), and stored at −20°C for at least 24 hours before imaging.

Vessel structure was assessed using the same protocol, substituting anti-AQP4 (Abcam, ab259318; 1:250) and anti-CD13 (Abcam, ab33489; 1:250) in the primary step, and lectin-DL488 (Vector, DL1174; 1:200) with appropriate secondary antibodies (Jackson AB 2632470 and Invitrogen A11077; 1:250) in the secondary step. Slides were mounted with ProLong Diamond Antifade with DAPI (Invitrogen, P36962).

Iron deposits were identified by Prussian blue staining.^18^ Slides were immersed for 20 minutes in a 1:1 mixture of 20% hydrochloric acid and 10% potassium ferrocyanide (Sigma-Aldrich, #P3289), washed three times in distilled water, and counterstained with nuclear fast red (ThermoFisher, #211980050) for 5 minutes. Slides were dehydrated through graded alcohols, cleared in xylene, and mounted with Permount (Fisher Chemical, SP15).

### Slide imaging and analysis

All slides were digitized using an Olympus VS200 scanning microscope at 20x magnification. All imaging and image analysis were performed by operators blinded to treatment.

NeuN-positive area was quantified by automated pixel classification in QuPath (v0.5.1).^19^ ROIs were manually drawn in the hippocampus and cortex; pixel classification thresholds were determined automatically using Otsu’s method and applied uniformly. Results were expressed as the percentage of positive pixels per ROI, averaged across 1–3 sections per mouse.

Prussian blue-positive area was quantified in Fiji (v.2.14.0/1.54f).^17^ Positively-stained areas were isolated by color thresholding (hue 111–196, saturation 13–255, brightness 0–255), and the size measured using the Analyze Particles function. Values were averaged across 2–3 sections per mouse.

Lectin, CD13, and AQP4 were quantified by automated pixel classification in QuPath. Two 500 µm × 500 µm ellipse annotations were drawn per hemisphere in the cortex. A pixel classifier was trained on 10–20 manually annotated positive and negative regions per image (ANN_MLP classifier; full resolution; Gaussian, Laplacian of Gaussian, and weighted deviation; scales 0.5 and 1.0), then applied to all four annotations with minimum object and hole sizes of 20 µm² and 500 µm², respectively. CD13 and AQP4 were quantified in the same ROIs using automated thresholding (Gaussian prefilter, sigma 2.5; thresholds of 600 and 4000, respectively). Results were expressed as detections per mm², averaged across 1–3 sections per mouse.

### Statistical analysis

All analyses were performed in Prism (v.10.4.1, GraphPad). Group differences were assessed by Mann-Whitney test unless otherwise noted. Data are presented as mean ± SEM, with *p* < 0.05 considered statistically significant. All operators were blinded to treatment group throughout.

## RESULTS

### Whole-brain irradiation reduces pial arteriolar flow without altering cortical tissue perfusion

To determine whether WBI produces lasting reductions in cerebral perfusion, we measured resting cortical blood flow by LSCI one year after irradiation. Global mean blur rate (MBR) across the imaging field was significantly reduced in WBI-treated mice compared to controls (1028 vs. 1199, *p* < 0.05; **Fig. 1A, D**). To resolve the spatial distribution of this effect, we segmented the imaging field into pial vessel and cortical tissue compartments using an MBR threshold of 890.9. The fractional area occupied by pial flow signal was markedly reduced in WBI-treated mice (69% vs. 87%, *p* < 0.05; **Fig. 1B, E**), and mean MBR within the pial compartment was significantly decreased (1154 vs. 1261, *p* < 0.05; **Fig. 1F**). By contrast, MBR in the cortical tissue compartment was unchanged (743 vs. 738, *p* > 0.05; **Fig. 1C, G**). The concurrent reduction in pial flow area and pial flow velocity, without a corresponding change in cortical tissue perfusion, indicates that WBI produces a persistent upstream reduction in pial arteriolar flow rather than a global decrease in parenchymal perfusion.

**Figure 1:**
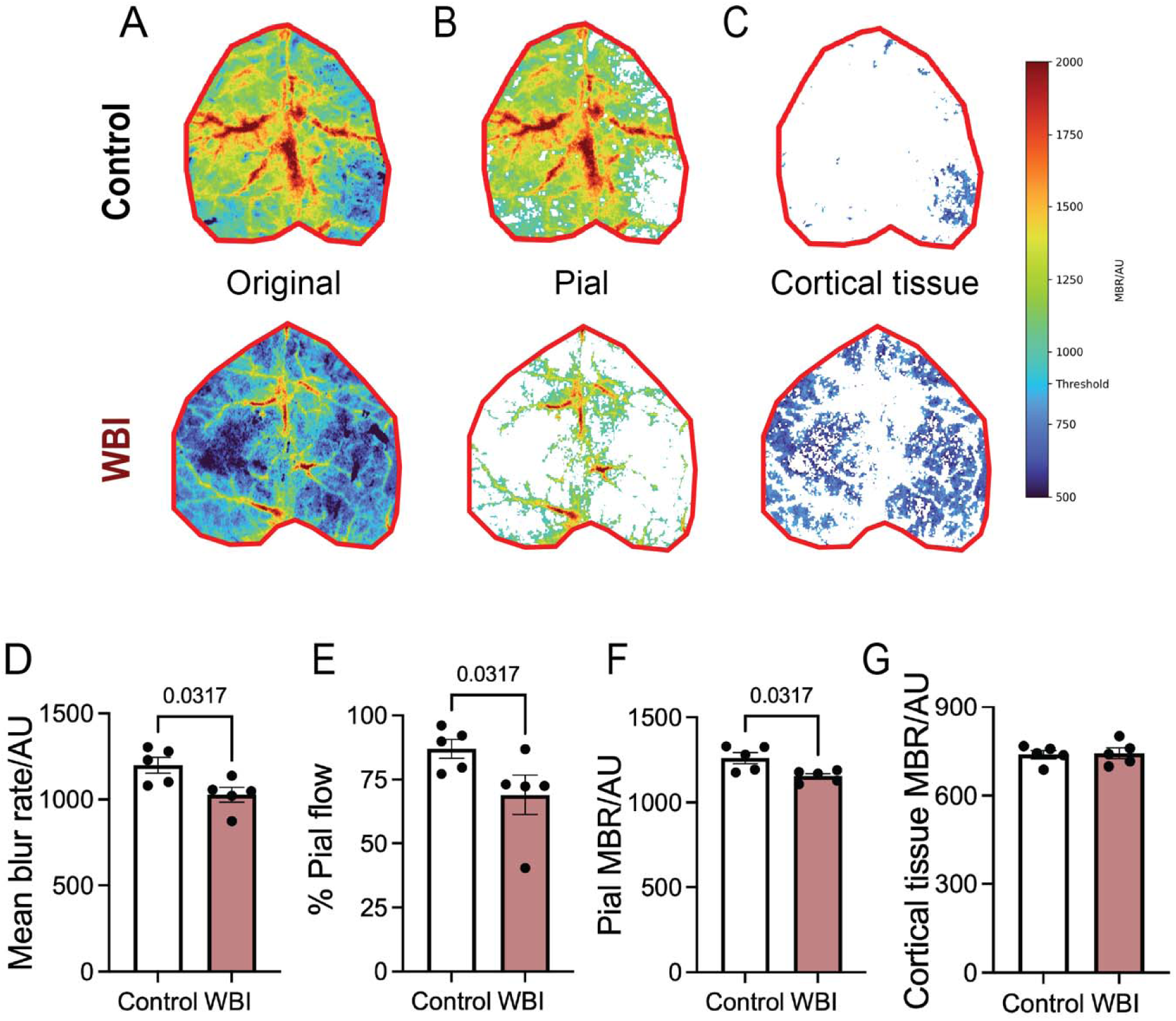
Whole-brain irradiation reduces pial flow velocity and area fraction without altering cortical tissue perfusion. **(A)** Representative overall blood flow, measured as mean blur rate (MBR), in control (top) and WBI-treated (bottom) mice. A threshold of MBR 891 was applied to segment pial **(B)** and cortical tissue **(C)** flow. **(D)** Global blood flow, before thresholding. **(E)** Fraction of the analyzed area occupied by pial flow signal. **(F, G)** Average blood flow in the pial (F) and cortical tissue (G) segments. Statistical analyses were performed by Mann-Whitney test (D-G). Exact *p* values indicated when *p* < 0.05. For (D-G), *n* = 5 mice per group.

### Whole-brain irradiation induces cerebral atrophy and anxiety-like behavior

To characterize the structural consequences of late-stage RIBI, we quantified regional brain volumes by atlas-based MRI segmentation and neuronal density by NeuN immunostaining. WBI-treated mice exhibited significant volume reductions in whole brain and total grey matter (*p* < 0.05; **Fig. 2C–D**), as well as in multiple regions implicated in anxiety and stress physiology, including the amygdala, brainstem, cerebellum, cerebrum, hypothalamus, and periaqueductal gray (all *p* < 0.05; **Fig. 2F–I, L–M**). The total white matter, frontal cortex, and hippocampus compartments of WBI-treated mice were not significantly affected by WBI, nor were there any changes in the number of NeuN-positive nuclei in the cortex or hippocampus (*p* > 0.05; **Fig. 2E, J, K, N-Q**). However, because neuronal density was assessed as a proportion of tissue area rather than as absolute cell counts, these data do not exclude the possibility that total neuron number declined proportionally with tissue volume. Stereological quantification would be required to resolve this distinction. Nevertheless, the absence of any change in relative neuronal density indicates that irradiation did not produce disproportionate neuronal loss beyond that which could be accounted for by overall tissue atrophy.

**Figure 2:**
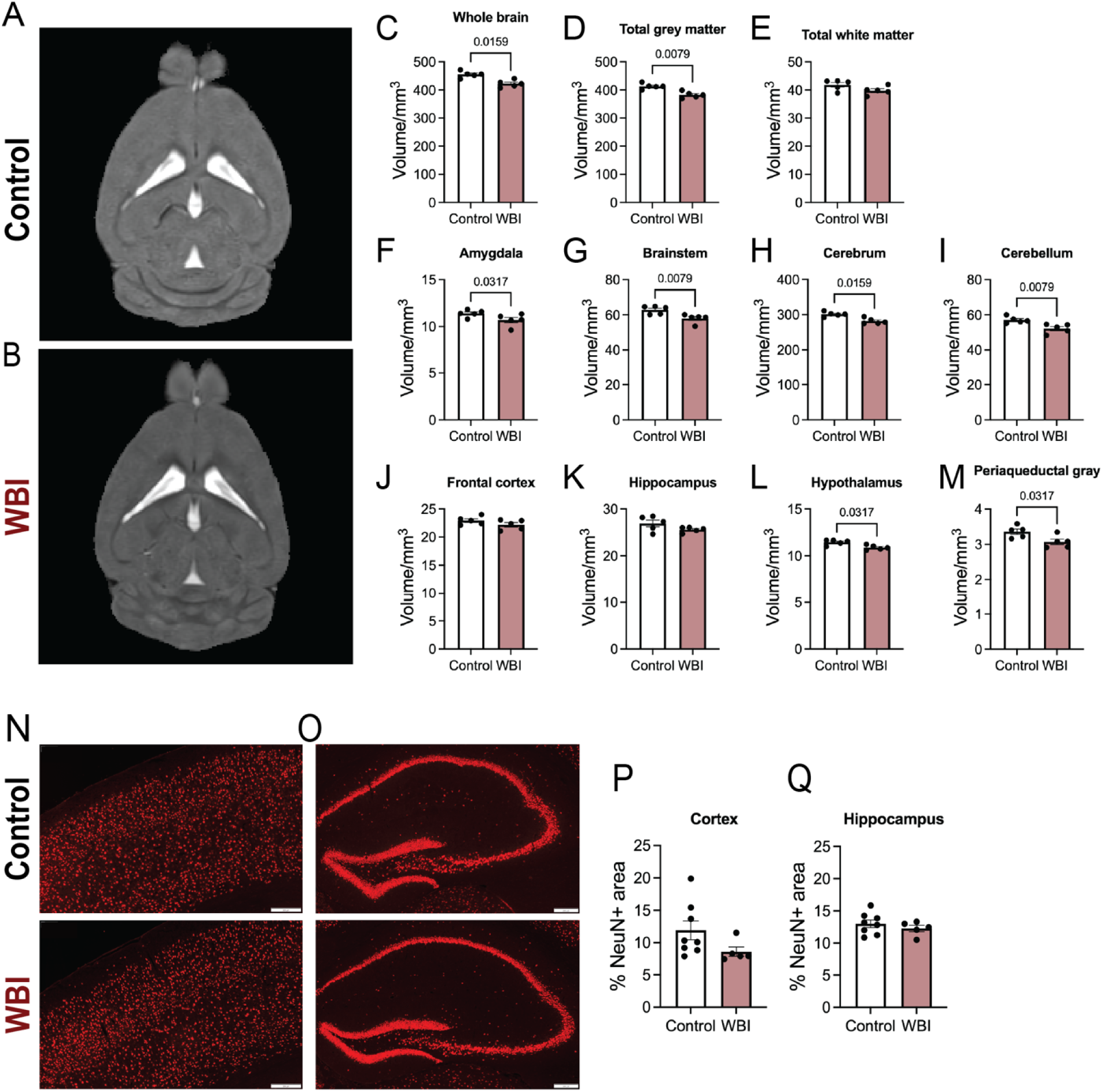
Whole-brain irradiation induces cerebral atrophy without loss of neuronal nuclei. **(A-B)** Representative T2-weighted anatomical MRI scans from a control (A) and WBI-treated (B) mouse. **(C-M)** Tissue volume was measured in the whole brain (C), grey matter (D), white matter (E), amygdala (F), brainstem (G), cerebrum (H), cerebellum (I), frontal cortex (J), hippocampus (K), hypothalamus (L), and periaqueductal gray (M) compartments, as defined by the DSURQE atlas. **(N-O)** Representative NeuN-positive stained images for neuronal nuclei in the cortex (N) and hippocampus (O) of control (top) and WBI-treated (bottom) mice. Scale bar indicates 200 μm. **(P-Q)** NeuN-positive area as a percentage of total tissue area in the cortex (P) and hippocampus (Q) of control and WBI-treated mice. Statistical analyses were performed by Mann-Whitney test (C-M, P-Q). Exact *p* values indicated when *p* < 0.05. For (C-M), *n* = 5 mice per group; for (P-Q), *n* = 8 Control, 5 WBI-treated mice.

Behaviorally, WBI-treated mice displayed significantly increased thigmotaxis in the open field test (86% vs. 74% time in border zone, *p* < 0.01; **Fig. 3A–B**), consistent with heightened anxiety-like behavior. Total distance traveled and mean velocity were unchanged (*p* > 0.05; **Fig. 3C–D**), excluding reduced locomotor activity as a confound. Novelty preference and discrimination index in the Y-maze did not differ between groups (*p* > 0.05; **Fig. 3E–G**). Together with preserved hippocampal volume and relative neuronal density, these findings indicate that the behavioral phenotype at one year is dominated by anxiety rather than gross hippocampal structural disruption.

**Figure 3:**
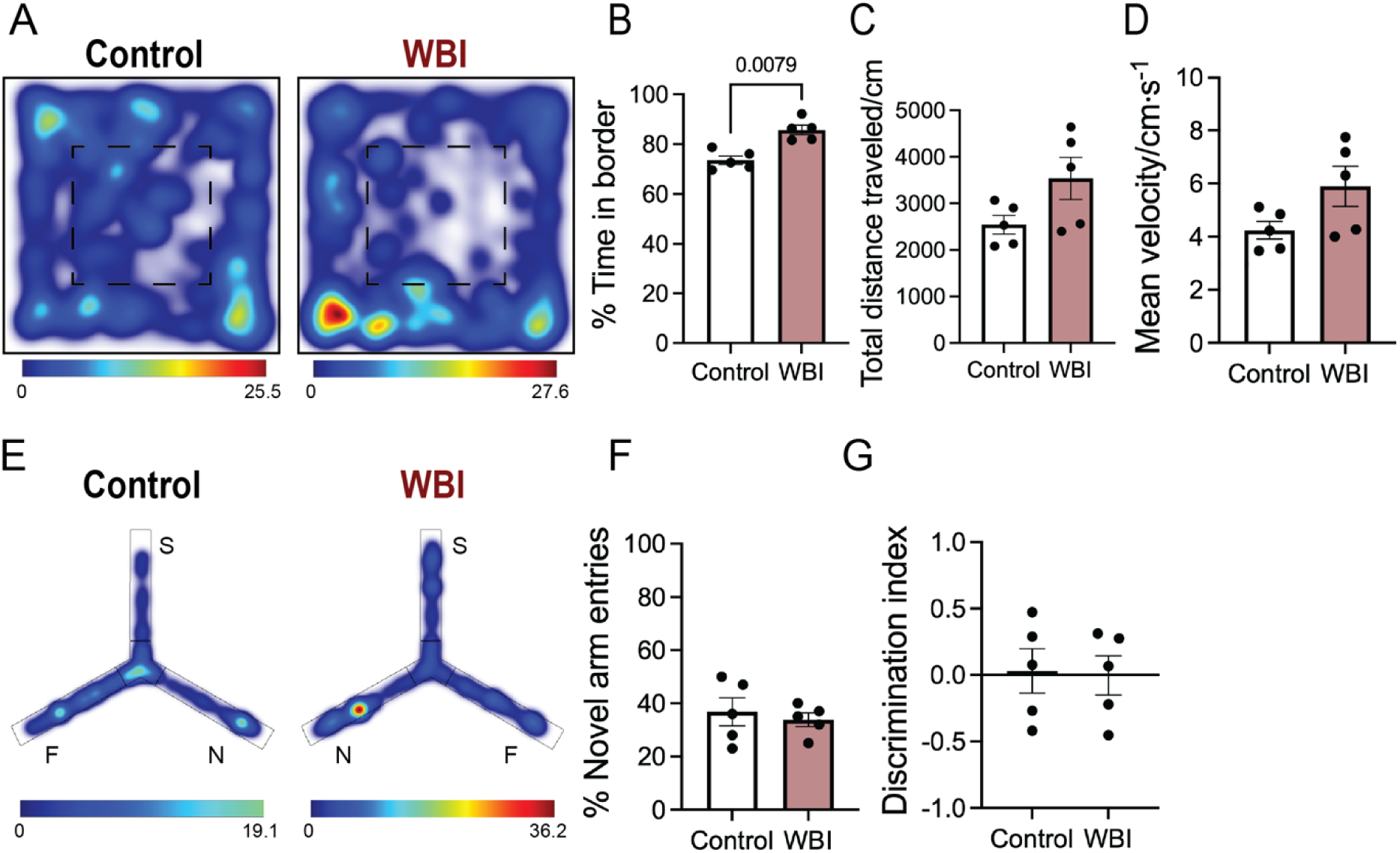
Whole-brain irradiation increases anxiety-like behavior without impairing spatial working memory. **(A**) Representative heat maps indicating the amount of time spent in different areas of the open field arena for control (left) and WBI-treated (right) mice. The dashed square indicates the center zone; the remaining area represents the border zone. **(B)** Percentage time spent in the border zones. **(C, D)** Locomotor function was evaluated by total distance traveled (C) and mean velocity (D) during the open field test. **(E)** Representative heat maps indicating the amount of time spent in the different areas of the Y-maze arena for control (left) and WBI-treated (right) mice during the novel arm phase. Mice were tested 4 hours after first exposure to the start (S) and familiar (F) arms, with preference for novelty based on time spent in the novel (N) arm. **(F, G)** Percentage of novel arm entries (F) and discrimination index (G). Color scales in (A, E) measure seconds. Statistical analyses were performed by Mann-Whitney test (B-D, F-G). Exact *p* values indicated when *p* < 0.05. For (B-D, F-G), *n* = 5 mice per group.

### Whole-brain irradiation induces tissue iron deposition and blood-brain barrier disruption consistent with chronic microhemorrhage

To assess whether pial hypoperfusion was accompanied by broader cerebrovascular injury, we evaluated vascular integrity across the brain using T2*-weighted MRI, Prussian blue histology, and molecular markers of the neurovascular unit. T2* relaxation time - an index of paramagnetic iron content - was significantly shortened in the whole brain, total grey matter, cerebrum, and hypothalamus of WBI-treated mice (all *p* < 0.01; **Fig. 4C–D, H, L**), with a trend toward reduction in the frontal cortex (*p* = 0.056; **Fig. 4J**). These data suggest increased iron content. Regions including the amygdala, brainstem, cerebellum, hippocampus, and periaqueductal gray showed no significant change (all *p* > 0.05; **Fig. 4E–G, I–K, M**). Prussian blue staining revealed a significant increase in hemosiderin-positive deposits in WBI-treated mice (mean area per section 84.5 vs. 767 µm², *p* < 0.05; **Fig. 4N–P**), corroborating that the T2* signal changes reflect iron accumulation from prior red blood cell extravasation and chronic microhemorrhage. Notably, the regional distribution of T2* changes extended well beyond the superficial cortex assessed by LSCI, providing convergent evidence that cerebrovascular injury in this model is not limited to pial vessels.

**Figure 4:**
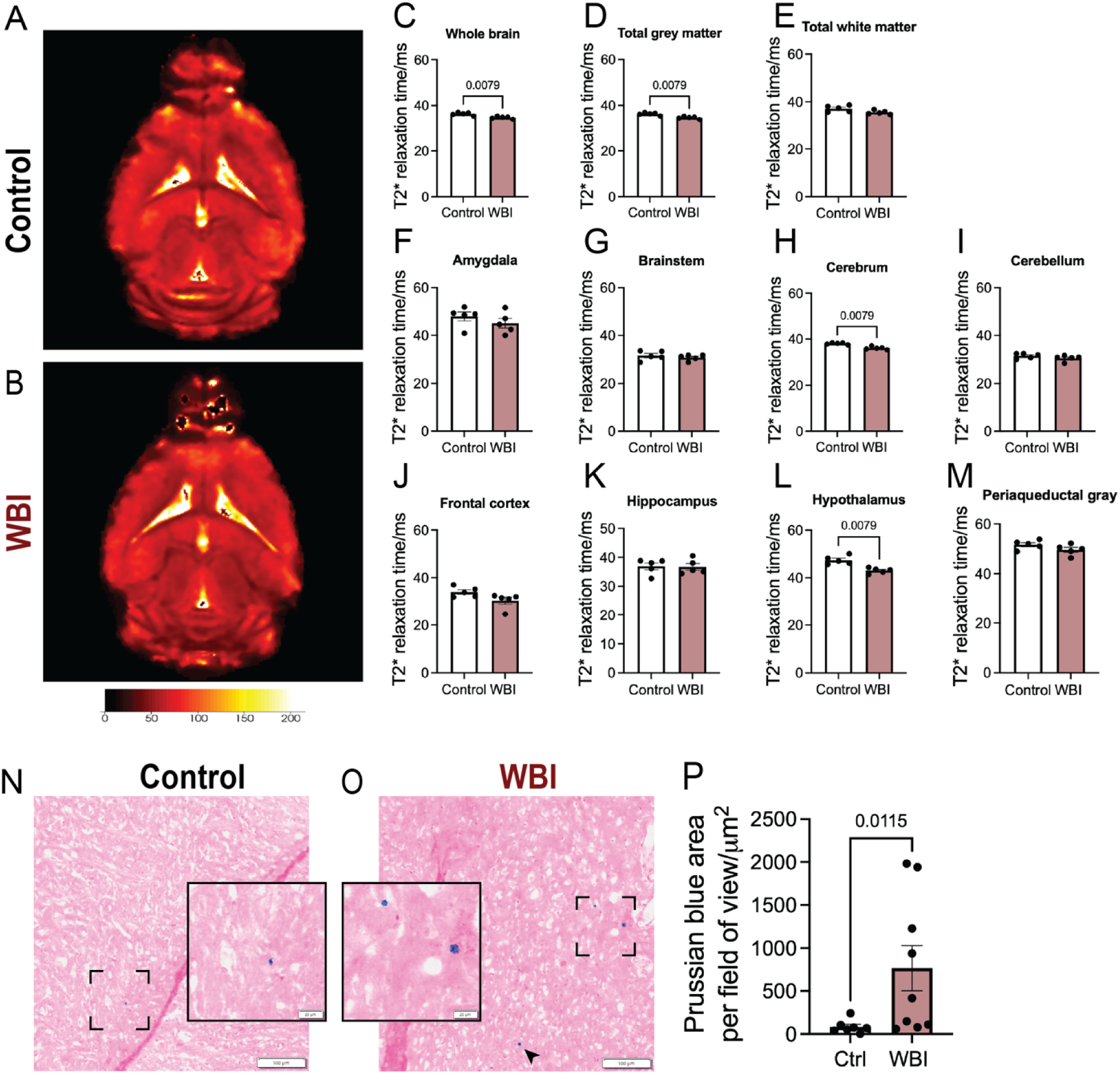
Whole-brain irradiation induces tissue iron deposition consistent with chronic microhemorrhage. **(A,B)** Representative T2*-weighted MRI images from a control (A) and WBI-treated (B) mouse. **(C-M)** T2* relaxation time was measured in the whole brain (C), grey matter (D), white matter (E), amygdala (F), brainstem (G), cerebrum (H), cerebellum (I), frontal cortex (J), hippocampus (K), hypothalamus (L), and periaqueductal gray (M) compartments, as defined by the DSURQE atlas. **(N-O)** Representative Prussian blue-stained images for hemosiderin deposits consistent with microhemorrhages of control (N) and WBI-treated (O) mice. Insets show magnified regions of Prussian blue-positive staining highlight the difference in staining between control and WBI-treated mice. Scale bar indicates 100 μm for the main image and 20 μm for the subset. Black arrow in (O) indicates a Prussian blue-positive deposit outside of the inset area. **(P)** Total Prussian blue-positive area captured per field of view. Statistical analyses were performed by Mann-Whitney test (C-M, P). Exact *p* values indicated when *p* < 0.05. For (C-M), *n* = 5 mice per group; for (P), *n* = 7 Control, 9 WBI-treated mice.

At the molecular level, immunoblot of isolated cerebral microvessels revealed significantly reduced expression of ZO-1 and VE-cadherin in WBI-treated mice (*p* < 0.05; **Fig. 5A–C**), indicating loss of endothelial tight junction and adherens junction integrity essential for blood-brain barrier function. Despite this molecular evidence of barrier disruption, structural markers of neurovascular unit composition were preserved relative to tissue area: lectin-positive vessel density, CD13-positive pericyte coverage, and AQP4-positive astrocyte endfeet were all unchanged in the cortex of WBI-treated mice (all *p* > 0.05; **Fig. 5D–G**). As with neuronal density, these vascular density measures are expressed relative to tissue area and thus cannot exclude proportional loss of vascular elements concurrent with brain atrophy. Nonetheless, blood-brain barrier disruption and pial hypoperfusion were observed without detectable changes in the relative density of any neurovascular unit component, implicating functional endothelial impairment rather than overt structural cellular dropout as the primary pathological substrate of late RIBI.

**Figure 5:**
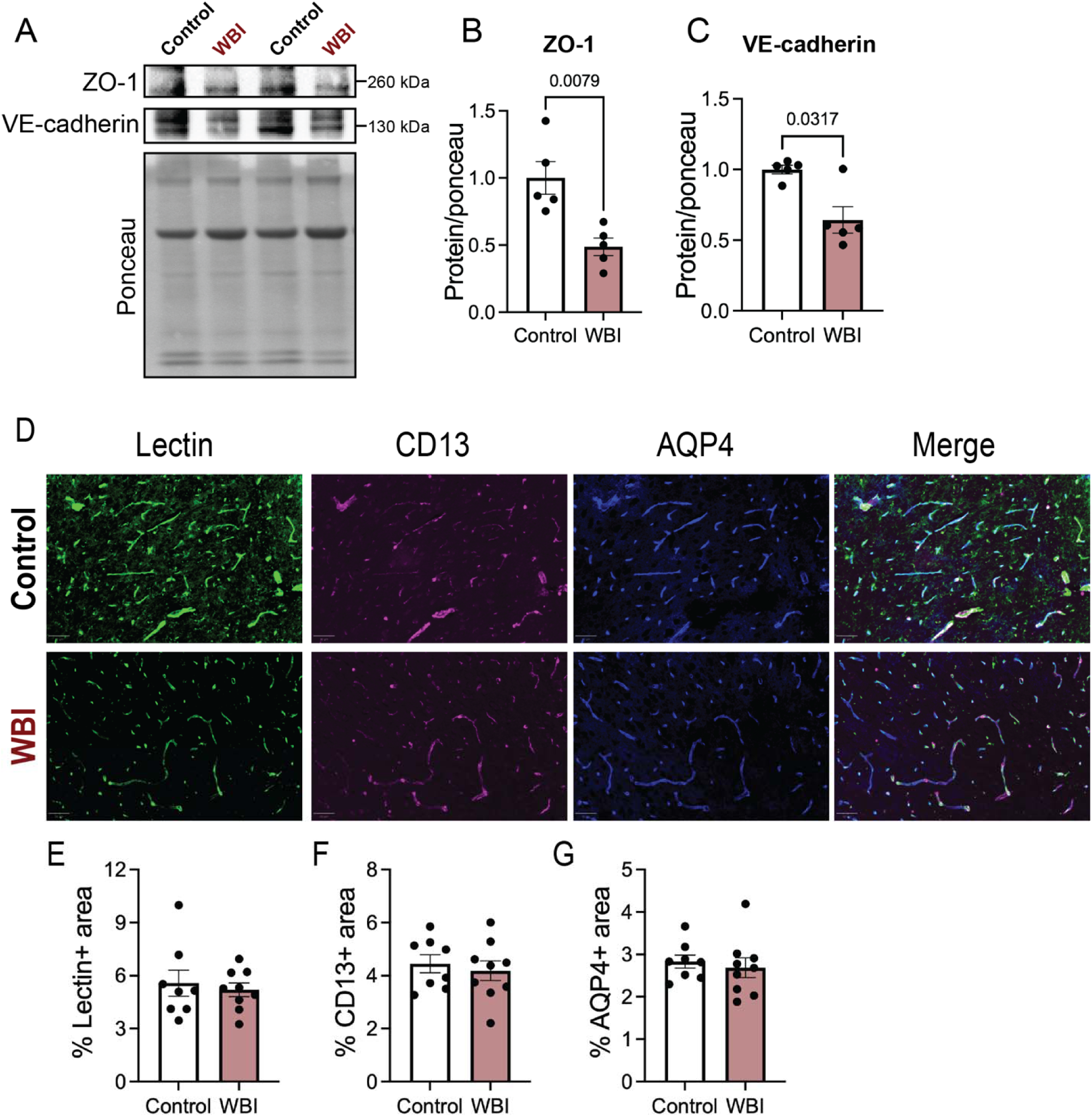
Whole-brain irradiation disrupts the blood-brain barrier without overt loss of capillaries, pericytes, or astrocyte endfeet. **(A)** Immunoblot of ZO-1 and VE-cadherin expression in isolated microvessels of representative control and WBI-treated mice, with corresponding Ponceau S total protein staining as loading control. **(B, C)** Quantification of ZO-1 (B) and VE-cadherin (C) expression as represented in (A). **(D)** Representative lectin, CD13, AQP4, and composite images from the cortex of a control (top) and WBI-treated (bottom) mouse. Scale bars indicate 50 μm. **(E-G)** Quantification of the percent lectin- (E), CD13- (F), and AQP4-positive (G) staining as in (D). Statistical analyses were performed by Mann-Whitney test (B-C, E-G). Exact *p* values indicated when *p* < 0.05. For (B-C), *n* = 5 mice per group; for (E-G), *n* = 8 Control, 9 WBI-treated mice.

## DISCUSSION

Cranial radiotherapy produces lasting damage to the cerebral vasculature, yet the relative contributions of structural vessel loss versus functional vascular impairment to the chronic sequelae of RIBI remain poorly defined. Here, using a comprehensive multimodal approach in adult mice at one year following clinically-relevant 12 Gy WBI, we show that late-stage RIBI is characterized by a distinct pattern of pial arteriolar dysfunction coexisting with widespread parenchymal microvascular injury, in the absence of detectable rarefaction of the cerebral vasculature or disproportionate neuronal loss. Specifically, WBI reduced pial arteriolar flow area and velocity - measured as a direct index of upstream cerebrovascular function by LSCI - while cortical tissue perfusion measured in the same imaging field was preserved. This pial hypoperfusion was accompanied by a spatially broader pattern of parenchymal injury evidenced by multi-regional cerebral atrophy, tissue iron deposition and microhemorrhage on MRI and histology, and molecular evidence of active endothelial barrier leakage reflected by loss of ZO-1 and VE-cadherin in isolated microvessels. That these vascular and structural changes unfolded against a backdrop of preserved relative vascular and neuronal density underscores the central conclusion of this work: that functional endothelial impairment, operating across multiple levels of the cerebrovascular hierarchy, is sufficient to drive the chronic pathological phenotype of late RIBI without the structural vessel loss that has historically dominated mechanistic discussions of this condition.

The use of resting perfusion as the primary vascular outcome distinguishes this study from prior LSCI investigations in WBI models, which focused on stimulus-evoked blood flow responses and demonstrated early impairment of neurovascular coupling and endothelial reactivity.^9,10^ Resting perfusion reflects the baseline functional state of the cerebral vasculature and may more sensitively capture chronic endothelial dysfunction than evoked responses alone. The dissociation we observed between pial flow reduction and preserved cortical tissue perfusion is itself informative: it suggests that the primary site of vascular dysfunction in this model lies at the level of the pial arterioles, which regulate downstream cortical perfusion, rather than at the capillary bed. Whether this pial dysfunction eventually propagates to impair parenchymal perfusion at later timepoints, or whether cortical tissue perfusion is maintained through autoregulatory compensation, remains an important open question. This pattern is consistent with our group’s prior clinical observation of reduced resting retinal perfusion detectable within months of ocular radiotherapy - preceding capillary dropout and vision loss by years^11^ - and suggests that pial arteriolar dysfunction may represent an early and sensitive indicator of radiation-induced vascular injury more broadly.

Notably, pial hypoperfusion and barrier disruption occurred without detectable changes in the relative density of lectin-positive vessels, CD13-positive pericytes, or AQP4-positive astrocyte endfeet, consistent with the hypothesis that functional impairment of surviving vascular cells, rather than their loss, drives the cerebrovascular phenotype of late RIBI. This is in keeping with the broader literature using moderate, clinically-relevant radiation doses, where a recent systematic review of 72 preclinical studies found preserved vascular structure alongside cognitive and vascular impairments.^2^ A recent study using 15 Gy WBI similarly reported pericyte senescence and cognitive impairment without changes in cellular density,^20^ and our group’s proteomic analysis of isolated cerebral microvessels at 6 months post-12 Gy WBI identified significant dysregulation of mitochondrial and metabolic proteins.^21^ Together, these data converge on a model in which radiation induces a state of endothelial and pericyte dysfunction - potentially driven by senescence, metabolic impairment, or oxidative stress - that reduces pial arteriolar flow and compromises barrier integrity in the absence of capillary rarefaction. It is important to note, however, that vascular density measures in the present study are expressed relative to tissue area; given the significant brain atrophy observed, proportional vascular loss concurrent with tissue volume reduction cannot be formally excluded without absolute volumetric quantification.

The observed cerebral atrophy and preserved relative neuronal density warrant careful interpretation. Grey matter volume was significantly reduced across multiple regions, yet NeuN-positive neuronal density, expressed as a proportion of tissue area, was unchanged in both cortex and hippocampus. As discussed in the Results, this finding does not exclude proportional neuronal loss concurrent with atrophy, and stereological quantification of absolute neuron number would be required to resolve this question definitively. An alternative explanation is that atrophy reflects loss of neuropil - particularly dendritic arbor, which has been shown to be reduced by up to 70% after cranial irradiation^22^ - rather than neuronal dropout. This interpretation is consistent with the predominantly grey matter distribution of volume loss observed by MRI, given that grey matter volume is composed largely of dendrites, axons, and extracellular matrix. The age of irradiation may also be relevant: studies using adult rather than juvenile rodents have demonstrated cognitive impairment without loss of mature neurons or reduction in neurogenesis,^23^ suggesting that mechanisms other than neuronal loss account for functional deficits in the adult-irradiated brain.

Behaviorally, irradiated mice exhibited increased anxiety-like behavior in the open field test without impairment of spatial working memory in the Y-maze. The anxiety phenotype is consistent with the pattern of structural changes observed (significant atrophy of the amygdala, hypothalamus, and periaqueductal gray, all structures integral to fear and stress responses), while preserved hippocampal volume and relative neuronal density align with the absence of Y-maze deficits. These findings suggest that at one year post-12 Gy WBI in adult mice, the behavioral consequences of RIBI are regionally selective, manifesting as anxiety-like behavior in the context of preserved hippocampal-dependent memory. Whether memory deficits emerge at later timepoints, or at higher doses, remains to be determined.

Several limitations of this study merit acknowledgment. First, all assessments were performed at a single late timepoint, precluding conclusions about the temporal sequence of vascular dysfunction, barrier disruption, atrophy, and behavioral change. Longitudinal studies with serial LSCI and histological assessment are needed to resolve whether pial hypoperfusion precedes or follows barrier disruption and iron accumulation, and whether capillary rarefaction occurs transiently at earlier timepoints before resolving. Second, LSCI measures perfusion in superficial pial and cortical vessels and does not provide direct information about deeper parenchymal or white matter perfusion; the MRI data provide complementary but indirect evidence of broader vascular injury. Third, as noted above, density measures for both neuronal and vascular elements are expressed relative to tissue area, and absolute quantification by stereology would be needed to exclude proportional cellular loss. Fourth, the decreased T2* relaxation time may reflect increased deoxyhemoglobin secondary to hypoperfusion in addition to tissue iron accumulation, though the convergent Prussian blue and LSCI findings independently support both pathologies. Finally, this study used male mice only; given known sex differences in radiation sensitivity and cerebrovascular physiology, whether these findings generalize to female animals requires investigation.

Despite these limitations, the co-occurrence of pial hypoperfusion, blood-brain barrier disruption, and widespread tissue iron accumulation in the absence of detectable structural vessel or disproportionate neuronal loss points to functional endothelial impairment as a primary and therapeutically actionable mechanism of late RIBI.

## Author contributions

Conceptualization, MRT, MSP, IMG; experimental procedures, MRT, AHV, PL, AC, VS, LHL; analysis, MRT, AHV, CYL, EDA; writing - original draft, MRT, IMG; writing - review and editing, MRT, AHV, PL, AC, CYL, EDA, VS, LHL, WT, MSP, IMG; supervision, WT, MSP, IMG.

## Ethics statement

All experiments were conducted with the approval of the University of Iowa and the Iowa City VA Health Care System Institutional Animal Care and Use Committees.

## Data availability

All data supporting the findings of this study are available from the corresponding author upon reasonable request.

## Acknowledgements

The authors acknowledge the technical services of the University of Iowa Orthopedics Histopathology Service Center for their assistance with slide imaging and Prussian blue staining. The authors also acknowledge the Ionizing Radiation Services Facility in the Department of Radiation Oncology at the University of Iowa.

