## Supplemental figure 1 for "Late radiation-induced brain injury is associated with functional cerebrovascular impairment without structural vessel loss in mice"

**Running title:**  Hypoperfusion without rarefaction in late RIBI

**Corresponding author:**
Isabella M. Grumbach MD, PhD

University of Oklahoma

Department of Internal Medicine

6444 AAT

800 Stanton Young Blvd

Oklahoma City, OK 73104, USA

405-271-8001 ext. 55435

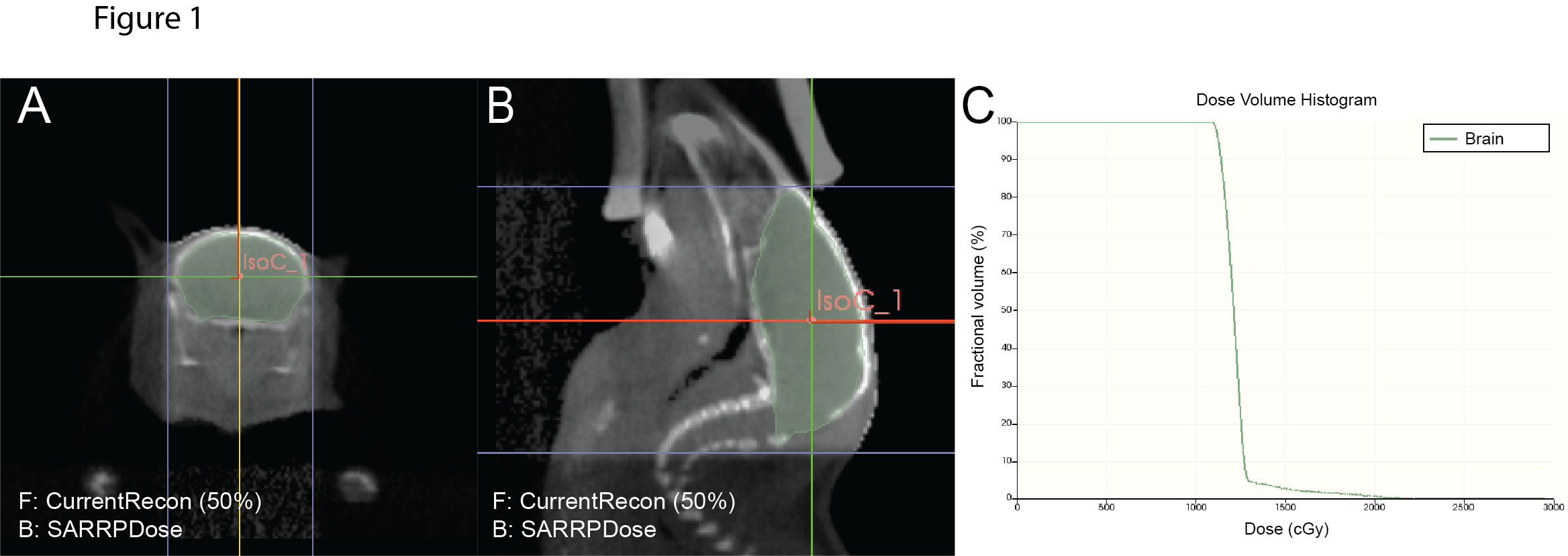

**Supplementary Figure 1: Treatment plan for whole-brain irradiation.** Representative axial (A) and sagittal (B) views with the whole brain volume contoured in green, and isocenter indicated by salmon-colored IsoC_1. A prescribed dose of 12 Gy was delivered to the isocenter. (C) Dose delivery was confirmed by dose-volume histogram of the contoured brain.
